# PRISMA: A precision functional imaging dataset of autistic and non-autistic adults

**DOI:** 10.64898/2026.01.12.698952

**Authors:** Nadza Dzinalija, Hilde M. Geurts, H. Steven Scholte, Joe Bathelt

## Abstract

Autism spectrum conditions are characterised by substantial heterogeneity in brain function, yet most neuroimaging studies rely on brief single-session acquisitions that prioritise group-level comparisons over individual characterisation. We present a precision functional imaging dataset with extended neuroimaging acquisitions to advance research on individual differences in autism. The dataset comprises 33 autistic adults between the ages of 18 and 30, and 34 age- and gender-matched comparison participants, each scanned across three sessions with up to 90 minutes of functional magnetic resonance imaging per participant. Functional data were acquired during eyes-open rest, news viewing, and reality television viewing to probe state-dependent differences in network engagement. The dataset includes high-resolution structural imaging, diffusion-weighted imaging with dual phase-encoding, and physiological recordings of cardiac and respiratory activity acquired simultaneously with functional scans. All data are organised following Brain Imaging Data Structure principles and accompanied by comprehensive quality control metrics and preprocessed derivatives. This dataset enables investigation of individual-specific functional brain organisation, test-retest reliability of connectivity patterns, and state-dependency of functional networks in autism, addressing a notable gap in publicly available neuroimaging resources.

## Background & Summary

Autism spectrum disorder is characterised by pronounced heterogeneity in clinical presentation, cognitive profiles, and underlying neurobiology. Over the past two decades, functional magnetic resonance imaging (fMRI) has emerged as a primary tool for investigating the neural basis of autism, with early studies typically examining small, independent samples of 20-40 participants (Philip et al., 2012). Whilst these studies provided initial insights into atypical functional connectivity patterns, particularly within the default mode network and sensory-processing regions, their limited statistical power and varied methodological approaches hindered the identification of replicable findings.

The establishment of large-scale data-sharing initiatives, most notably the Autism Brain Imaging Data Exchange (ABIDE) (Di Martino et al., 2012, 2017), represented a paradigm shift in the field. By aggregating resting-state fMRI data from over 2,000 participants across multiple sites, ABIDE enabled researchers to conduct well-powered investigations of functional connectivity differences in autism. Subsequent mega- and meta-analyses of these pooled datasets have confirmed the presence of widespread connectivity alterations (Ilioska et al., 2023), yet have simultaneously revealed substantial heterogeneity both within the autism population and across individual studies (Benkarim et al., 2021; Dickie et al., 2018). This heterogeneity suggests that group-level averaging may obscure clinically meaningful individual variation.

Concurrently, a complementary line of research in human neuroscience has demonstrated that individual differences in functional brain organisation can be reliably characterised when sufficient data are collected from each participant (Demeter & Greene, 2024). The Midnight

Scan Club and similar precision imaging studies have shown that extended scanning protocols can capture stable, person-specific features of functional connectivity that remain obscured in brief, single-session acquisitions (Demeter & Greene, 2024; Gordon et al., 2017; Gratton et al., 2020; Laumann et al., 2015). These “connectome fingerprints” not only enable reliable identification of individuals across scanning sessions but also show stronger associations with behavioural phenotypes than group-averaged connectivity patterns (Ramduny & Kelly, 2024). Critically, recent evidence suggests that autistic individuals may exhibit greater idiosyncrasy in their functional brain organisation compared to neurotypical controls (Benkarim et al., 2021), potentially explaining some of the heterogeneity observed in large-scale studies.

Despite these methodological advances, no publicly available dataset has yet combined precision imaging approaches with the study of autism. Existing datasets either provide brief acquisitions from large samples (e.g., ABIDE, EU-AIMS LEAP) or extended acquisitions from neurotypical individuals only (e.g., Midnight Scan Club, Human Connectome Project). This represents a notable gap in available resources, as understanding individual-level functional brain organisation in autism may be crucial for developing personalised intervention strategies and for resolving the heterogeneity that has complicated clinical translation.

Here, we present a precision functional imaging dataset of 33 autistic adults and 34 age-and sex-matched neurotypical comparison participants, each scanned across three separate sessions with up to 90 minutes of functional data per participant. To our knowledge, this represents the first openly shared dataset combining extended functional acquisitions with autism research. The dataset includes multiple viewing conditions (eyes-open rest, reality television clips, and news programme segments) designed to probe state-dependent differences in network engagement, as naturalistic viewing paradigms have been shown to elicit more stable individual differences and may better reveal connectivity patterns relevant to real-world social and cognitive processing (Finn & Bandettini, 2021; Vanderwal et al., 2017).

The dataset further includes high-resolution structural imaging (0.8mm isotropic T1-weighted and T2-weighted for improved cortical surface reconstruction) and diffusion-weighted imaging (128 directions, b=1000 s/mm², dual phase-encoding) to enable comprehensive characterisation of individual neuroanatomy and white matter connectivity. All functional data were acquired using multiband acceleration and processed with NORDIC denoising, ensuring high temporal signal-to-noise ratio (Moser et al., 2024). Further, we recorded heart rate and breathing during the scanning sessions for enhanced removal of physiological confounds (Lynch et al., 2020) that may be particularly problematic in autistic individuals (Cheng et al., 2020).

Sessions were separated by at least 5 days, allowing estimation of within-person stability versus between-session variability in functional connectivity patterns (Cho et al., 2021). Participants also reported on their thought content after each scan using a standardised questionnaire (Diaz et al., 2013; Simpraga et al., 2021) to characterise the association between connectivity patterns and spontaneous thought (Finn, 2021). Further, participants completed a behavioural battery assessing autistic traits, general cognition, and general psychological functioning, providing phenotypic data for sample characterisation and brain-behaviour analyses.

This dataset offers several opportunities for reuse. First, it enables investigation of individual-specific functional brain organisation in autism, including the stability and state-dependency of connectivity patterns. Second, the combination of functional, structural, and diffusion data permits multimodal characterisation of brain organisation, which may reveal structure-function relationships that are obscured in unimodal analyses. Third, the naturalistic viewing paradigms provide ecologically valid data that may better capture real-world processing differences. Finally, the inclusion of phenotypic data supports investigation of brain-behaviour relationships, especially when combining the data with larger-scale datasets e.g., through normative modelling (Marquand et al., 2016, 2019) or data fusion (Webb-Vargas et al., 2017). We anticipate that this dataset will be valuable for researchers developing methods for precision psychiatry, investigating neural heterogeneity in autism, and advancing individualised approaches to understanding neurodevelopmental conditions.

## Methods

### Participants

Participants were recruited through local advertisements and by contacting autism interest groups in the region. All participants received compensation for travel expenses and were reimbursed for their time at the standard institutional rate (€15 per hour).

Participants first completed a screening survey to determine eligibility. Inclusion criteria required that participants be native Dutch speakers, aged between 18 and 30 years, have normal or corrected-to-normal vision, and report no contraindications for MRI (e.g., medical implants, accidents involving metal, pregnancy). For inclusion in the autism group, participants were required to confirm a formal diagnosis of an autism spectrum condition by a medical professional or licensed psychologist, and to obtain a score of 6 or higher on the 10-item version of the Autism Spectrum Quotient (AQ-10) (Allison et al., 2012). Participants in the non-autistic comparison group were required to confirm that neither they nor any immediate family members (parents or siblings) had received an autism diagnosis, and had to reach an AQ-10 score below 6. Eligible individuals were subsequently contacted by telephone to confirm the absence of additional MRI contraindications and to schedule their scanning appointment.

One participant withdrew from the study after the first scanning session. All remaining participants completed the study.

### MRI Data Acquisition

MRI data were acquired between March 2024 and July 2025 using a Philips Achieva dStream 3T scanner (Philips, Best, The Netherlands). At the start of each scan session, a low-resolution survey scan was made, which was used to determine the location of the field-of-view.

High-resolution structural images were acquired using a three-dimensional T1-weighted turbo field echo (3DT1 TFE) sequence (TR = 9.9 ms, TE = 4.58 ms, flip angle = 8°, field of view [FoV] = 256 × 256 mm², 312 slices, voxel size = 0.8 × 0.8 × 0.8 mm³). In addition, a three-dimensional T2-weighted turbo spin echo (3DT2 TSE) sequence was collected (TR = 2500 ms, TE = 330 ms, flip angle = 90°, FoV = 240 × 256 mm², 312 slices, voxel size = 0.8 × 0.8 × 0.8 mm³).

Diffusion-weighted imaging data were acquired using a spin-echo EPI sequence (TR = 2529 ms, TE = 75.44 ms, 2 mm isotropic voxels, 63 slices). We acquired 128 diffusion-encoding directions at b = 1000 s/mm² with two b = 0 volumes, using reversed phase-encoding (AP/PA) acquisitions for distortion correction. Parallel imaging (SENSE = 1.3, multiband = 3) and partial Fourier (0.632) were employed to reduce scan time.

Blood-oxygen-level-dependent (BOLD) fMRI data were acquired using a gradient-echo echo-planar imaging (GE-EPI) sequence (TR = 1600 ms, TE = 29.93 ms, flip angle = 74°) with transverse slice orientation. Multiband acceleration (factor = 4) was applied with SENSE parallel imaging (factor = 1.5). The acquisition covered 60 slices (15 groups of 4) with an FoV of 224 × 224 × 131 mm³, voxel size = 2 × 2 × 2 mm³, and inter-slice gap = 0.2 mm. To enable susceptibility distortion correction, four additional volumes with reversed phase-encoding polarity were acquired at the start of each functional run (topup field maps, anterior–posterior).

During functional MRI scans, additional physiological data was recorded. Respiratory traces were recorded using a respiratory belt (air filled cushion) bound on top of the subject’s diaphragm using a Velcro band. Cardiac traces were recorded using a plethysmograph attached to the subject’s left index finger. Data was transferred to the scanner PC as plain-text files using a wireless recorder with a sampling frequency of 496 Hz.

#### Precision-Imaging Protocol

Participants underwent MRI scanning on three separate occasions, with a minimum interval of five days between successive sessions. The mean interval between sessions one and two in the autism group was 10.68 days (SD = 7.17 days) and in the comparison group was 9.91 days (SD = 5.07 days). Between sessions two and three the mean interval in the autism group was 10.77 days (SD = 7.10 days) and in the comparison group was 8.91 days (SD = 4.64 days). To facilitate acclimatisation to the scanner environment, structural imaging was performed at the start of each session. During the anatomical scans, participants watched a cartoon movie, which also served to verify that they could clearly see the projection screen and hear the audio. T1-weighted images were acquired during the first session, T2-weighted images during the second, and diffusion-weighted images during the third. If the initial T1-weighted acquisition was compromised by motion artefacts, a repeat acquisition was performed during the second session.

Following the structural sequence, participants completed three functional MRI runs of approximately 10 minutes each: (i) an eyes-open resting-state condition with a white fixation cross presented on a grey background, (ii) a video clip of a news programme, and (iii) a video clip of a reality-television show. All video clips were obtained from the official YouTube channel of the Dutch national broadcaster (NPO). The news clips were selected to present factual content with minimal emotional arousal, covering topics such as pro- and anti-European arguments in Poland, the economic drivers of farmer protests in the Netherlands, and cultural perspectives on monarchy in European countries. The reality-television clips, in contrast, were chosen for their emphasis on social interaction, and included a blind date between two individuals, a travel episode featuring three friends, and interviews with one or two participants speaking directly to the camera. For this condition, the first eight minutes of an episode featuring autistic individuals discussing common preconceptions (e.g. “autistic people are great at maths”) were combined with two minutes from the beginning of a programme on transgender people to achieve a total duration of 10 minutes. The video clips used are available with this dataset on OpenNeuro.org.

After each functional run, participants completed the Amsterdam Resting-State Questionnaire (ARSQ-2.0) (Diaz et al., 2013). Items were presented in randomised order on the screen. Participants rated the extent to which each statement reflected their experience on a five-point Likert scale, using three buttons on an MR-compatible response pad to move and confirm the marker position.

A scanning assistant communicated with participants between sequences to monitor their comfort and wellbeing. For autistic participants, the same scanning assistant accompanied each individual across all three scanning occasions to provide continuity and reduce variability.

### Data standardisation, preprocessing, and derivatives

#### Raw data standardisation

Raw imaging data were acquired in native Philips PAR/REC format and underwent a standardised conversion prior to organisation into the Brain Imaging Data Structure (BIDS) format. The data were first converted from PAR/REC to NIfTI format using dicm2nii v2023.03.16 (https://github.com/xiangruili/dicm2nii) implemented in MATLAB R2024b. This tool preserves essential header information including acquisition parameters and geometric orientation whilst generating NIfTI files compatible with downstream processing pipelines. Functional MRI and TOP-UP images for distortion correction underwent thermal noise reduction using NORDIC (NOise Reduction with DIstribution Corrected PCA) (Dowdle et al., 2023). NORDIC exploits the complex-valued nature of MRI data by utilising both magnitude and phase information to distinguish thermal noise from signal components via principal component analysis. This denoising step was implemented in MATLAB R2024b.

Following format conversion and denoising, data were renamed and organised according to BIDS specification (version 1.10.1) using in-house software. This process involved systematic renaming of files according to BIDS naming conventions and generation of associated metadata files (JSON sidecars, TSV files) containing acquisition parameters and participant information. To protect participant privacy, all structural images were defaced using MiDeFace 2, which is part of FreeSurfer v7.4.1 Docker image. The de-facing mask was derived using the T1-weighted images and applied to the T2-weighted images after corregistration using the ‘mri_coreg’ function of FreeSurfer. The scripts used for data processing are available via the GitHub repository (see “Code Availability” section for details). The BIDS validator v2.1.1 was used to check for errors or inconsistencies in the data organisation.

#### Data quality assessment

Quality control of anatomical and functional MRI data was performed using MRIQC v25.0.0 (Esteban et al., 2017), which generates standardised image quality metrics and visual reports for individual participant assessment. For diffusion-weighted imaging, data quality was evaluated using the quality control outputs generated by QSIPrep v1.0.2 (Cieslak et al., 2021).

#### Preprocessing of anatomical and functional MRI data

Results included in this manuscript come from preprocessing performed using the Docker version of fMRIPrep 25.2.3 (RRID:SCR_016216) (Esteban et al., 2019, 2025), which is based on Nipype 1.10.0 (RRID:SCR_002502) (Gorgolewski et al., 2011; Gorgolewski et al., 2017). The following boilerplate text was automatically generated by fMRIPrep and adapted for general description of the data processing with minimal changes.

##### Preprocessing of B0 inhomogeneity mappings

Fieldmaps were associated with the corresponding functional run for each participant. A B0-nonuniformity map (or fieldmap) was estimated based on two (or more) echo-planar imaging (EPI) references with topup (FSL) (Andersson et al., 2003).

##### Anatomical data preprocessing

T1-weighted (T1w) images were selected for each participant from the BIDS data. The T1w image was corrected for intensity non-uniformity (INU) with N4BiasFieldCorrection, distributed with ANTs 2.6.2 (RRID:SCR_004757) (Avants et al., 2008). The T1w-reference was then skull-stripped with a Nipype implementation of the antsBrainExtraction.sh workflow (from ANTs), using OASIS30ANTs as target template. Brain tissue segmentation of cerebrospinal fluid (CSF), white-matter (WM) and gray-matter (GM) was performed on the brain-extracted T1w using fast (FSL, RRID:SCR_002823) (Avants et al., 2008; Y. Zhang et al., 2001). Brain surfaces were reconstructed using recon-all (FreeSurfer 7.3.2, RRID:SCR_001847) (Dale et al., 1999). A T2-weighted image was used to improve pial surface refinement. Brain surfaces were reconstructed using recon-all (FreeSurfer 7.3.2, RRID:SCR_001847) (Dale et al., 1999), and the brain mask estimated previously was refined with a custom variation of the method to reconcile ANTs-derived and FreeSurfer-derived segmentations of the cortical gray-matter of Mindboggle (RRID:SCR_002438) (Klein et al., 2017). Volume-based spatial normalization to two standard spaces (MNI152NLin2009cAsym, MNI152NLin6Asym) was performed through nonlinear registration with antsRegistration (ANTs 2.6.2), using brain-extracted versions of both T1w reference and the T1w template. The following templates were selected for spatial normalization and accessed with TemplateFlow (25.0.4) (Ciric et al., 2022): ICBM 152 Nonlinear Asymmetrical template version 2009c (RRID:SCR_008796) (Fonov et al., 2009); TemplateFlow ID: MNI152NLin2009cAsym, FSL’s MNI ICBM 152 non-linear 6th Generation Asymmetric Average Brain Stereotaxic Registration Model (RRID:SCR_002823) (Evans et al., 2012); TemplateFlow ID: MNI152NLin6Asym.

##### Functional data preprocessing

For each of the up to 9 BOLD runs per subject (across all tasks and sessions), the following preprocessing was performed. First, a reference volume was generated, using a custom methodology of fMRIPrep, for use in head motion correction. Head-motion parameters with respect to the BOLD reference (transformation matrices, and six corresponding rotation and translation parameters) are estimated before any spatiotemporal filtering using mcflirt (FSL) (Jenkinson et al., 2002). The estimated fieldmap was then aligned with rigid-registration to the target EPI (echo-planar imaging) reference run. The field coefficients were mapped on to the reference EPI using the transform. The BOLD reference was then co-registered to the T1w reference using bbregister (FreeSurfer) which implements boundary-based registration (Greve & Fischl, 2009). Co-registration was configured with six degrees of freedom. The aligned T2w image was used for initial co-registration. Several confounding time-series were calculated based on the preprocessed BOLD: framewise displacement (FD), DVARS and three region-wise global signals. FD was computed using two formulations following Power (absolute sum of relative motions) (Power et al., 2014) and Jenkinson (relative root mean square displacement between affines) (Jenkinson et al., 2002). FD and DVARS are calculated for each functional run, both using their implementations in Nipype (following the definitions by Power et al. 2014)). The three global signals are extracted within the CSF, the WM, and the whole-brain masks. Additionally, a set of physiological regressors were extracted to allow for component-based noise correction (CompCor) (Behzadi et al., 2007). Principal components are estimated after high-pass filtering the preprocessed BOLD time-series (using a discrete cosine filter with 128s cut-off) for the two CompCor variants: temporal (tCompCor) and anatomical (aCompCor). tCompCor components are then calculated from the top 2% variable voxels within the brain mask. For aCompCor, three probabilistic masks (CSF, WM and combined CSF+WM) are generated in anatomical space. The implementation differs from that of Behzadi et al. in that instead of eroding the masks by 2 pixels on BOLD space, a mask of pixels that likely contain a volume fraction of GM is subtracted from the aCompCor masks. This mask is obtained by dilating a GM mask extracted from the FreeSurfer’s aseg segmentation, and it ensures components are not extracted from voxels containing a minimal fraction of GM. Finally, these masks are resampled into BOLD space and binarized by thresholding at 0.99 (as in the original implementation). Components are also calculated separately within the WM and CSF masks. For each CompCor decomposition, the k components with the largest singular values are retained, such that the retained components’ time series are sufficient to explain 50 percent of variance across the nuisance mask (CSF, WM, combined, or temporal). The remaining components are dropped from consideration. The head-motion estimates calculated in the correction step were also placed within the corresponding confounds file. The confound time series derived from head motion estimates and global signals were expanded with the inclusion of temporal derivatives and quadratic terms for each (Satterthwaite et al., 2013). Frames that exceeded a threshold of 0.5 mm FD or 1.5 standardized DVARS were annotated as motion outliers. Additional nuisance timeseries are calculated by means of principal components analysis of the signal found within a thin band (crown) of voxels around the edge of the brain, as proposed by (Patriat et al., 2017). The BOLD time-series were resampled onto the left/right-symmetric template “fsLR” using the Connectome Workbench (Glasser et al., 2013). Grayordinates files (Glasser et al., 2013) containing 91k samples were also generated with surface data transformed directly to fsLR space and subcortical data transformed to 2 mm resolution MNI152NLin6Asym space. All resamplings can be performed with a single interpolation step by composing all the pertinent transformations (i.e. head-motion transform matrices, susceptibility distortion correction when available, and co-registrations to anatomical and output spaces). Gridded (volumetric) resamplings were performed using nitransforms, configured with cubic B-spline interpolation. Grayordinate “dscalar” files containing 91k samples were resampled onto fsLR using the Connectome Workbench (Glasser et al., 2013).

Many internal operations of fMRIPrep use Nilearn 0.11.1 (RRID:SCR_001362) (Abraham et al., 2014), mostly within the functional processing workflow. For more details of the pipeline, see the section corresponding to workflows in fMRIPrep’s documentation (see https://fmriprep.readthedocs.io/en/latest/workflows.html).

#### Postprocessing of fMRIPrep output

The eXtensible Connectivity Pipeline- DCAN (XCP-D) (Ciric et al., 2018; Mehta et al., 2024; Satterthwaite et al., 2013) was used to post-process the outputs of fMRIPrep version 25.2.3 (RRID:SCR_016216) (Esteban et al., 2019, 2020). XCP-D was built with Nipype version 1.10.0 (RRID:SCR_002502) (Gorgolewski et al., 2011). The following description was automatically generated by XCP-D and was minimally adjusted to reflect the processing across the entire dataset.

##### Segmentations

The following atlases were used in the workflow: the Schaefer Supplemented with Subcortical Structures (4S) atlas (King et al., 2019; Najdenovska et al., 2018; Pauli et al., 2018; Schaefer et al., 2018) at 10 different resolutions (1056, 156, 256, 356, 456, 556, 656, 756, 856, and 956 parcels), the Glasser atlas (Glasser et al., 2016), the Gordon atlas (Gordon et al., 2016), the Tian subcortical atlas (Gordon et al., 2016; Tian et al., 2020), the HCP CIFTI subcortical atlas (Glasser et al., 2013), the MIDB precision brain atlas derived from ABCD data and thresholded at 75% probability (Hermosillo et al., 2024), and the Myers-Labonte infant atlas thresholded at 50% probability (Myers et al., 2023).

##### Anatomical data

Native-space T1w images were transformed to MNI152NLin2009cAsym space at 1 mm3 resolution.

##### Functional data

For each of the up to 9 BOLD runs per subject (across all tasks and sessions), the following post-processing was performed: In total, 36 nuisance regressors were selected from the preprocessing confounds, according to the ‘36P’ strategy. These nuisance regressors included six motion parameters, mean global signal, mean white matter signal, mean cerebrospinal fluid signal with their temporal derivatives, and quadratic expansion of six motion parameters, tissue signals and their temporal derivatives (Ciric et al., 2018; Satterthwaite et al., 2013). The BOLD data were converted to NIfTI format, despiked with AFNI’s 3dDespike, and converted back to CIFTI format.

Nuisance regressors were regressed from the BOLD data using a denoising method based on Nilearn’s approach. The timeseries were band-pass filtered using a(n) second-order Butterworth filter, in order to retain signals between 0.01-0.08 Hz. The same filter was applied to the confounds. The resulting time series were then denoised using linear regression. The denoised BOLD was then smoothed using Connectome Workbench with a Gaussian kernel (FWHM=6 mm).

The amplitude of low-frequency fluctuation (ALFF) (Zou et al., 2008) was computed by transforming the mean-centered, standard deviation-normalized, denoised BOLD time series to the frequency domain. The power spectrum was computed within the 0.01-0.08 Hz frequency band and the mean square root of the power spectrum was calculated at each voxel to yield voxel-wise ALFF measures. The resulting ALFF values were then multiplied by the standard deviation of the denoised BOLD time series to retain the original scaling. The ALFF maps were smoothed with the Connectome Workbench using a Gaussian kernel (FWHM=6 mm).

For each hemisphere, regional homogeneity (ReHo) (Jiang & Zuo, 2016) was computed using surface-based 2dReHo (Zhang et al., 2019). Specifically, for each vertex on the surface, the Kendall’s coefficient of concordance (KCC) was computed with nearest-neighbor vertices to yield ReHo. For the subcortical, volumetric data, ReHo was computed with neighborhood voxels using AFNI’s 3dReHo (Taylor & Saad, 2013).

Processed functional timeseries were extracted from residual BOLD using Connectome Workbench (Marcus et al., 2011) for the atlases. Corresponding pair-wise functional connectivity between all regions was computed for each atlas, which was operationalized as the Pearson’s correlation of each parcel’s unsmoothed timeseries with the Connectome Workbench. In cases of partial coverage, uncovered vertices (values of all zeros or NaNs) were either ignored (when the parcel had >50.0% coverage) or were set to zero (when the parcel had <50.0% coverage).

Many internal operations of XCP-D use AFNI (Cox, 1996; Cox & Hyde, 1997),Connectome Workbench (Marcus et al., 2011), ANTS (Avants & Tustison, 2009), TemplateFlow version 25.0.3 (Ciric et al., 2022), matplotlib version 3.10.5 (Hunter, 2007), Nibabel version 5.3.2 (Brett et al., 2025), Nilearn version 0.12.1 (Abraham et al., 2014), numpy version 2.2.6 (Harris et al., 2020), pybids version 0.19.0 (Yarkoni et al., 2019), and scipy version 1.15.3 (Virtanen et al., 2020). For more details, see the XCP-D website (https://xcp-d.readthedocs.io).

#### Preprocessing of diffusion-weighted images (DWI)

Preprocessing was performed using QSIPrep 1.0.2 (Cieslak et al., 2021), which is based on Nipype 1.9.1 (RRID:SCR_002502) (Gorgolewski et al., 2011; Gorgolewski et al., 2017).

The anatomical reference image was reoriented into AC-PC alignment via a 6-DOF transform extracted from a full Affine registration to the MNI152NLin2009cAsym template. A full nonlinear registration to the template from AC-PC space was estimated via symmetric nonlinear registration (SyN) using antsRegistration. Brain extraction was performed on the T1w image using SynthStrip (Hoopes et al., 2022) and automated segmentation was performed using SynthSeg (Billot et al., 2023) (@synthseg2) from FreeSurfer version 7.3.1.

##### Diffusion data preprocessing

Images were grouped into two phase encoding polarity groups. DWI data were denoised using the Marchenko-Pastur PCA method implemented in dwidenoise (Billot et al., 2023; Cordero-Grande et al., 2019; Tournier et al., 2019; Veraart et al., 2016) with an automatically-determined window size of 5 voxels. After MP-PCA, Gibbs ringing was removed using MRtrix3 (Kellner et al., 2016; Tournier et al., 2019). Any images with a b-value less than 50 s/mm^2 were treated as a b=0 image. The mean intensity of the DWI series was adjusted so all the mean intensity of the b=0 images matched across each separate DWI scanning sequence. B1 field inhomogeneity was corrected using dwibiascorrect from MRtrix3 with the N4 algorithm (Tustison et al., 2010) after corrected images were resampled. Both distortion groups were then merged into a single file, as required for the FSL workflows.

FSL’s eddy was used for head motion correction and Eddy current correction (Andersson et al., 2016). Eddy was configured with a q-space smoothing factor of 10, a total of 5 iterations, and 1000 voxels used to estimate hyperparameters. A quadratic first level model and a linear second level model were used to characterize Eddy current-related spatial distortion. q-space coordinates were forcefully assigned to shells. Field offset was attempted to be separated from subject movement. Shells were aligned post-eddy. Eddy’s outlier replacement was run (Andersson et al., 2016). Data were grouped by slice, only including values from slices determined to contain at least 250 intracerebral voxels. Groups deviating by more than 4 standard deviations from the prediction had their data replaced with imputed values.

Data was collected with reversed phase-encode blips, resulting in pairs of images with distortions going in opposite directions. FSL’s TOPUP (Andersson et al., 2003) was used to estimate a susceptibility-induced off-resonance field based on b=0 images extracted from multiple DWI series with reversed phase encoding directions. The TOPUP-estimated fieldmap was incorporated into the Eddy current and head motion correction interpolation. Final interpolation was performed using the jac method.

Several confounding time-series were calculated based on the preprocessed DWI: framewise displacement (FD) using the implementation in Nipype (following the definitions by Power et al., 2014). The head-motion estimates calculated in the correction step were also placed within the corresponding confounds file. Slicewise cross correlation was also calculated. The DWI time-series were resampled to ACPC, generating a preprocessed DWI run in ACPC space with 1.75mm isotropic voxels.

Many internal operations of QSIPrep use Nilearn 0.10.1 (RRID:SCR_001362) (Abraham et al., 2014) and Dipy (Garyfallidis et al., 2014). For more details of the pipeline, see the section corresponding to workflows in QSIPrep’s documentation (see https://qsiprep.readthedocs.io/en/latest/workflows.html).

#### Postprocessing of QSIPrep output

Reconstruction was performed using QSIRecon 1.1.2 (Cieslak et al., 2021), which is based on Nipype 1.9.1 (RRID:SCR_002502) (Cieslak et al., 2021; Esteban et al., 2025; K. Gorgolewski et al., 2011).

A hybrid surface/volume segmentation was created (Smith et al., 2020). FreeSurfer outputs were registered to the QSIRecon outputs.

##### Anatomical data for DWI reconstruction

T1w-based spatial normalization calculated during preprocessing was used to map atlases from template space into alignment with DWIs. Brain masks from antsBrainExtraction were used in all subsequent reconstruction steps. The following atlases were used in the workflow: the Schaefer Supplemented with Subcortical Structures (4S) atlas (Glasser et al., 2016; King et al., 2019; Najdenovska et al., 2018; Pauli et al., 2018; Schaefer et al., 2018; Smith et al., 2020) at 5 different resolutions (156, 256, 356, 456, and 556 parcels), the Brainnetome 246-parcel atlas (Fan et al., 2016) extended with subcortical parcels, and the Gordon 333-parcel atlas (Gordon et al., 2016) extended with subcortical parcels. Cortical parcellations were mapped from template space to DWIS using the T1w-based spatial normalization.

##### MRtrix3 Reconstruction

Multi-tissue fiber response functions were estimated using the dhollander algorithm. FODs were estimated via constrained spherical deconvolution (CSD) (Tournier et al., 2004, 2008) using an unsupervised multi-tissue method (Dhollander et al., 2016; Dhollander & Raffelt, 2019). A single-shell-optimized multi-tissue CSD was performed using MRtrix3Tissue (https://3Tissue.github.io), a fork of MRtrix3 (Tournier et al., 2019) FODs were intensity-normalized using mtnormalize (Raffelt et al., 2017).

Many internal operations of QSIRecon use Nilearn 0.10.1 (RRID:SCR_001362) (Abraham et al., 2014) and Dipy 1.10.0 (Garyfallidis et al., 2014). For more details of the pipeline, see the section corresponding to workflows in QSIRecon’s documentation (see https://qsirecon.readthedocs.io/en/latest/workflows.html).

### Behavioural Assessment

Participants completed the Matrix Reasoning subtest of the Wechsler Adult Intelligence Scale – Fourth Edition (WAIS-IV-NL) to assess non-verbal cognitive ability (Wechsler, 2008). To evaluate verbal ability, the first 9 participants completed the Peabody Picture Vocabulary Test (PPVT-NL) (Dunn & Dunn, 1965); however, due to observed ceiling effects, all subsequent participants completed the WAIS-IV Vocabulary subtest instead. Trained research assistants administered these standardised tests according to the manual instructions. All assessments were conducted in Dutch, the native language of all participants.

Participants also completed the 60-item Autism Spectrum Quotient (AQ-60-NL) to assess autism-related traits (Hoekstra et al., 2008), and the young adult version of the Strengths and Difficulties Questionnaire (SDQ-NL) to assess general behavioural and emotional functioning (Brann et al., 2018). Additionally, participants completed a bespoke questionnaire to collect demographic information and details pertaining to autism diagnosis (Geurts et al., 2021).

### Data Records

The PRISMA dataset is organised according to the Brain Imaging Data Structure (BIDS) specification version 1.10.1 and is publicly available at doi:10.18112/openneuro.ds007182.v1.0.0 under accession number ds007182. The dataset has been validated using the BIDS Validator v2.1.1 with no errors. All neuroimaging data are stored as compressed NIfTI files (.nii.gz) with accompanying JSON sidecar files containing acquisition parameters. Physiological recordings are stored as compressed tab-separated value files (.tsv.gz) following BIDS specification for physiological recordings. Behavioural and phenotypic data are provided as tab-separated value files (.tsv) with JSON metadata files. This dataset uses a modified organisation where functional data across all three sessions are stored in a single directory per participant. Session assignment for each scan is documented in ‘sub-<label>_scans.tsv’.

### Technical Validation

**Sample characteristics & behavioural data quality.** Demographic characteristics are summarised in Table 1. The autism and comparison groups were well-matched on age, sex, handedness, and years of education. Both groups comprised university students and scored in the above-average range on measures of general cognitive ability (WAIS matrix reasoning scaled score ASC: M = 10.55, SD = 3.56; CMP: M = 11.88, SD = 2.72). As expected, the autism group reported significantly higher autistic traits on the AQ-50 (see Table 2). The average score in the autism group fell below the clinical screening threshold of 145 proposed by Hoekstra et al. (Hoekstra et al. 2008), consistent with a university-based sample characterised by late diagnosis and high adaptive functioning. Participants in the autism group also scored higher on the SDQ, with higher scores for emotional symptoms, peer relationship problems, and lower scores for prosocial behaviour compared to the comparison group (see Table 2). ARSQ subscale scores showed adequate variance with no floor or ceiling effects in either group, and sum scores within the expected ranges using dimension definitions in both version 1.0 (Diaz et al., 2013) and version 2.0 (Diaz et al., 2014) (see Tables 3 and 4).

**Table 1.**
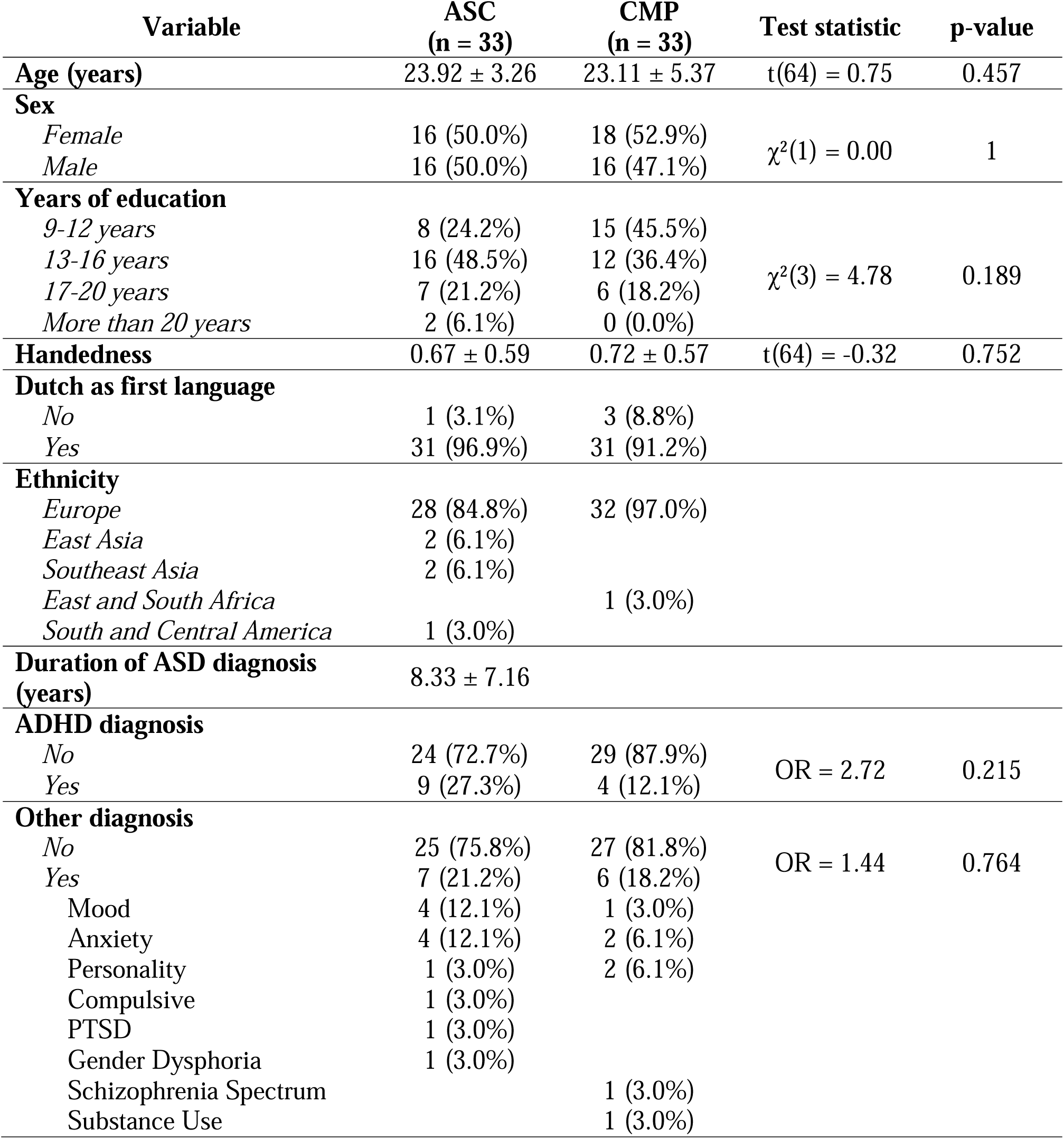
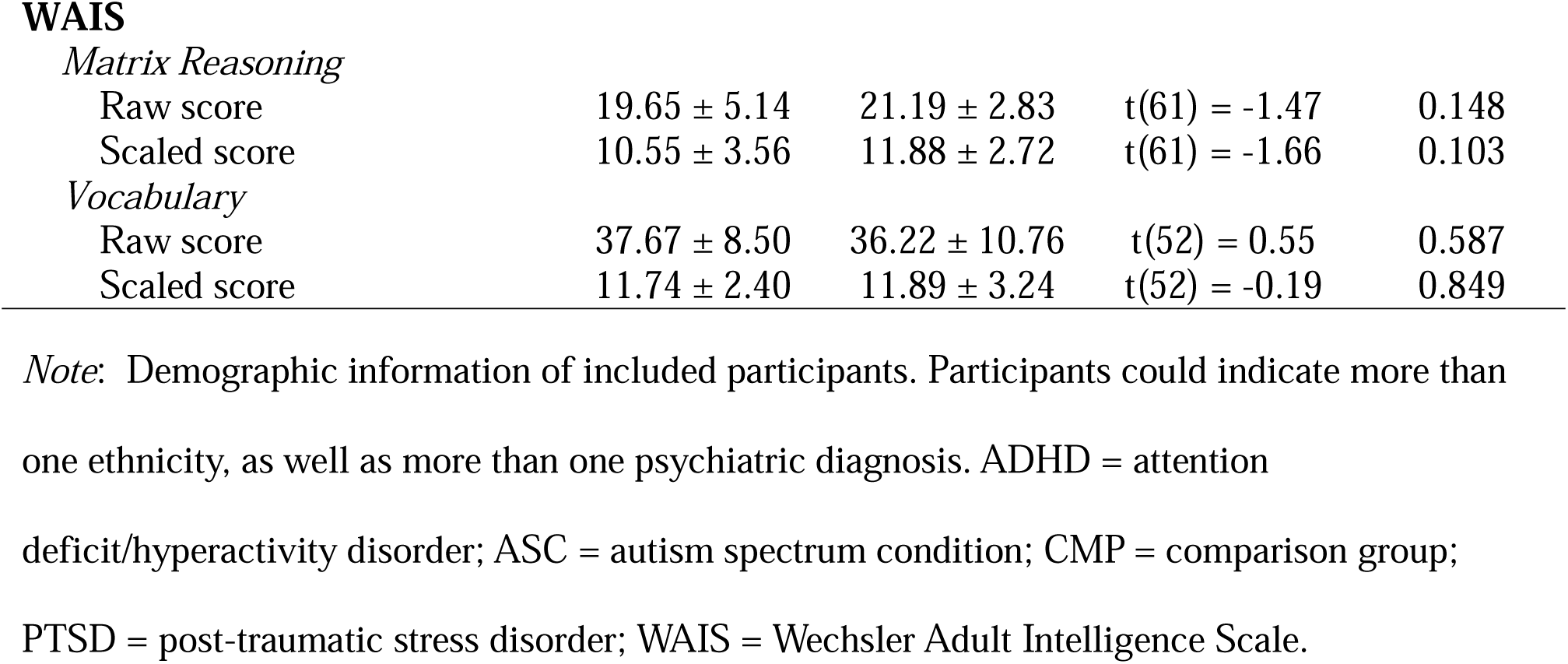
Demographic information of included participants.

| Variable | ASC<br>(n = 33) | CMP<br>(n = 33) | Test statistic | p-value |
| --- | --- | --- | --- | --- |
| **Age (years)** | 23.92 ± 3.26 | 23.11 ± 5.37 | t(64) = 0.75 | 0.457 |
| **Sex** |  |  |  |  |
| *Female* | 16 (50.0%) | 18 (52.9%) | $\chi^2(1) = 0.00$ | 1 |
| *Male* | 16 (50.0%) | 16 (47.1%) |  |  |
| **Years of education** |  |  |  |  |
| *9-12 years* | 8 (24.2%) | 15 (45.5%) | $\chi^2(3) = 4.78$ | 0.189 |
| *13-16 years* | 16 (48.5%) | 12 (36.4%) |  |  |
| *17-20 years* | 7 (21.2%) | 6 (18.2%) |  |  |
| *More than 20 years* | 2 (6.1%) | 0 (0.0%) |  |  |
| **Handedness** | 0.67 ± 0.59 | 0.72 ± 0.57 | t(64) = -0.32 | 0.752 |
| **Dutch as first language** |  |  |  |  |
| *No* | 1 (3.1%) | 3 (8.8%) |  |  |
| *Yes* | 31 (96.9%) | 31 (91.2%) |  |  |
| **Ethnicity** |  |  |  |  |
| *Europe* | 28 (84.8%) | 32 (97.0%) |  |  |
| *East Asia* | 2 (6.1%) |  |  |  |
| *Southeast Asia* | 2 (6.1%) |  |  |  |
| *East and South Africa* |  | 1 (3.0%) |  |  |
| *South and Central America* | 1 (3.0%) |  |  |  |
| **Duration of ASD diagnosis (years)** | 8.33 ± 7.16 |  |  |  |
| **ADHD diagnosis** |  |  |  |  |
| *No* | 24 (72.7%) | 29 (87.9%) | OR = 2.72 | 0.215 |
| *Yes* | 9 (27.3%) | 4 (12.1%) |  |  |
| **Other diagnosis** |  |  |  |  |
| *No* | 25 (75.8%) | 27 (81.8%) | OR = 1.44 | 0.764 |
| *Yes* | 7 (21.2%) | 6 (18.2%) |  |  |
| Mood | 4 (12.1%) | 1 (3.0%) |  |  |
| Anxiety | 4 (12.1%) | 2 (6.1%) |  |  |
| Personality | 1 (3.0%) | 2 (6.1%) |  |  |
| Compulsive | 1 (3.0%) |  |  |  |
| PTSD | 1 (3.0%) |  |  |  |
| Gender Dysphoria | 1 (3.0%) |  |  |  |
| Schizophrenia Spectrum |  | 1 (3.0%) |  |  |
| Substance Use |  | 1 (3.0%) |  |  |
| **WAIS***Matrix Reasoning* |  |  |  |  |
| Raw score | 19.65 ± 5.14 | 21.19 ± 2.83 | t(61) = -1.47 | 0.148 |
| Scaled score | 10.55 ± 3.56 | 11.88 ± 2.72 | t(61) = -1.66 | 0.103 |
| *Vocabulary* |  |  |  |  |
| Raw score | 37.67 ± 8.50 | 36.22 ± 10.76 | t(52) = 0.55 | 0.587 |
| Scaled score | 11.74 ± 2.40 | 11.89 ± 3.24 | t(52) = -0.19 | 0.849 |

*Note:* Demographic information of included participants. Participants could indicate more than
one ethnicity, as well as more than one psychiatric diagnosis. ADHD = attention
deficit/hyperactivity disorder; ASC = autism spectrum condition; CMP = comparison group;
PTSD = post-traumatic stress disorder; WAIS = Wechsler Adult Intelligence Scale.

**Table 2.**
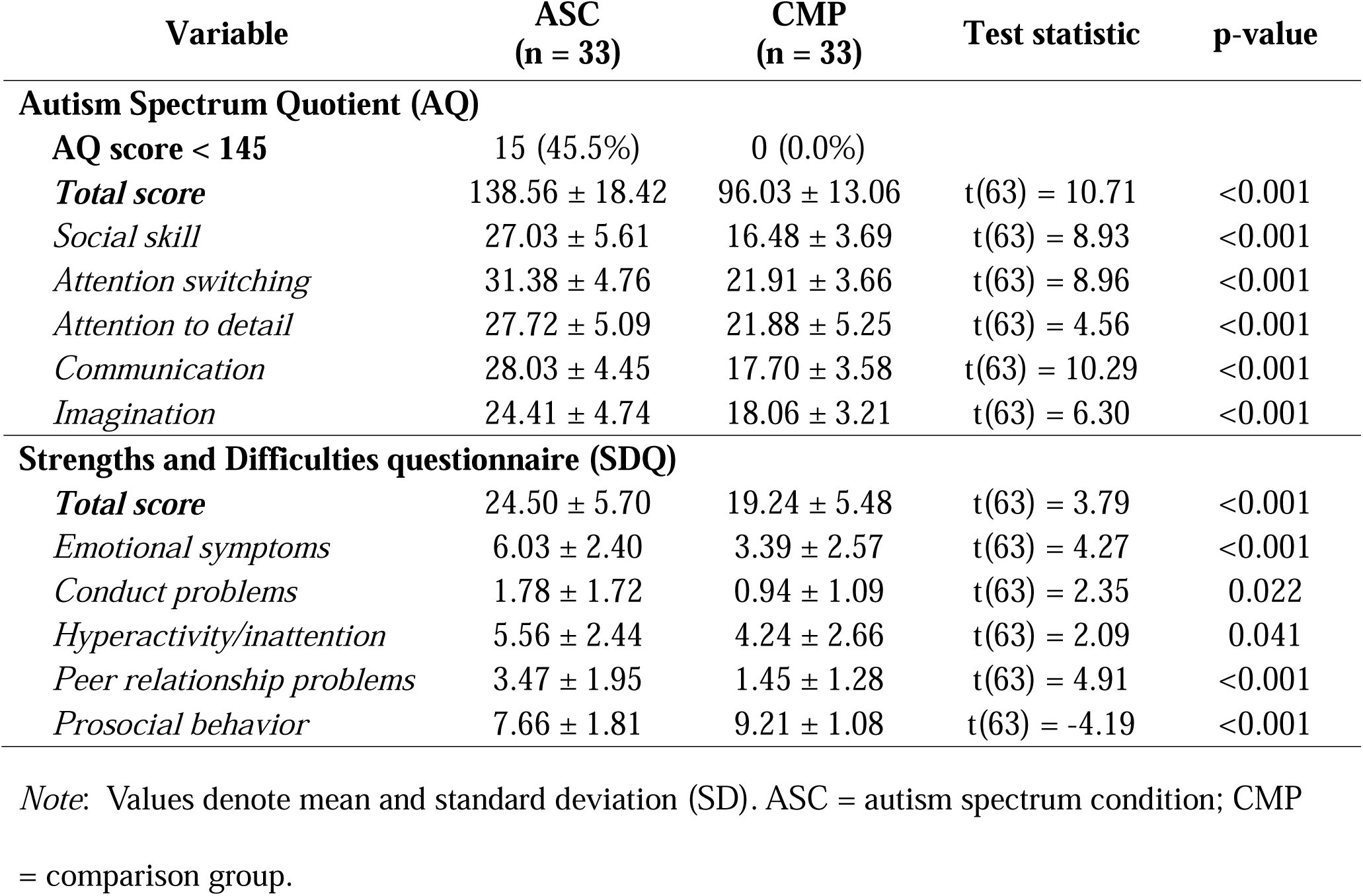
Descriptive statistics of autism-relevant behavioural and trait measures.

| Variable | ASC<br>(n = 33) | CMP<br>(n = 33) | Test statistic | p-value |
| --- | --- | --- | --- | --- |
| **Autism Spectrum Quotient (AQ)** |  |  |  |  |
| **AQ score < 145** | 15 (45.5%) | 0 (0.0%) |  |  |
| **Total score** | 138.56 ± 18.42 | 96.03 ± 13.06 | t(63) = 10.71 | <0.001 |
| *Social skill* | 27.03 ± 5.61 | 16.48 ± 3.69 | t(63) = 8.93 | <0.001 |
| *Attention switching* | 31.38 ± 4.76 | 21.91 ± 3.66 | t(63) = 8.96 | <0.001 |
| *Attention to detail* | 27.72 ± 5.09 | 21.88 ± 5.25 | t(63) = 4.56 | <0.001 |
| *Communication* | 28.03 ± 4.45 | 17.70 ± 3.58 | t(63) = 10.29 | <0.001 |
| *Imagination* | 24.41 ± 4.74 | 18.06 ± 3.21 | t(63) = 6.30 | <0.001 |
| **Strengths and Difficulties questionnaire (SDQ)** |  |  |  |  |
| **Total score** | 24.50 ± 5.70 | 19.24 ± 5.48 | t(63) = 3.79 | <0.001 |
| *Emotional symptoms* | 6.03 ± 2.40 | 3.39 ± 2.57 | t(63) = 4.27 | <0.001 |
| *Conduct problems* | 1.78 ± 1.72 | 0.94 ± 1.09 | t(63) = 2.35 | 0.022 |
| *Hyperactivity/inattention* | 5.56 ± 2.44 | 4.24 ± 2.66 | t(63) = 2.09 | 0.041 |
| *Peer relationship problems* | 3.47 ± 1.95 | 1.45 ± 1.28 | t(63) = 4.91 | <0.001 |
| *Prosocial behavior* | 7.66 ± 1.81 | 9.21 ± 1.08 | t(63) = -4.19 | <0.001 |

*Note:* Values denote mean and standard deviation (SD). ASC = autism spectrum condition; CMP = comparison group.

**Table 3.** Descriptive statistics of Amsterdam Resting State Questionnaire (ARSQ) version 1.0.

| **Video** | **ARSQ Domain Version 1.0** | **ASC** | **CMP** |
| --- | --- | --- | --- |
| News | Comfort | $9.54 \pm 1.72$ | $10.37 \pm 2.08$ |
| | Discontinuity of Mind | $14.73 \pm 1.87$ | $13.89 \pm 2.69$ |
| | Planning | $16.54 \pm 4.38$ | $15.73 \pm 3.60$ |
| | Self | $6.05 \pm 1.56$ | $5.68 \pm 1.46$ |
| | Sleep | $7.30 \pm 2.58$ | $6.89 \pm 2.62$ |
| | Somatic Awareness | $6.38 \pm 2.43$ | $6.96 \pm 2.47$ |
| | Theory of Mind | $8.15 \pm 2.27$ | $8.84 \pm 2.00$ |
| | Validation | $14.11 \pm 2.40$ | $13.55 \pm 1.53$ |
| Reality TV | Comfort | $10.21 \pm 1.91$ | $11.04 \pm 1.62$ |
| | Discontinuity of Mind | $14.33 \pm 2.20$ | $13.07 \pm 2.53$ |
| | Planning | $15.30 \pm 4.44$ | $13.17 \pm 3.68$ |
| | Self | $6.86 \pm 1.54$ | $5.97 \pm 1.50$ |
| | Sleep | $6.41 \pm 2.14$ | $5.74 \pm 1.80$ |
| | Somatic Awareness | $6.48 \pm 2.15$ | $6.40 \pm 2.77$ |
| | Theory of Mind | $9.91 \pm 2.29$ | $9.97 \pm 1.99$ |
| | Validation | $14.24 \pm 2.08$ | $13.49 \pm 1.84$ |
| Rest | Comfort | $9.54 \pm 2.36$ | $10.51 \pm 1.84$ |
| | Discontinuity of Mind | $17.08 \pm 2.28$ | $15.27 \pm 2.63$ |
| | Planning | $18.86 \pm 4.45$ | $20.27 \pm 4.06$ |
| | Self | $7.26 \pm 1.27$ | $7.07 \pm 1.23$ |
| | Sleep | $9.57 \pm 3.01$ | $9.22 \pm 3.03$ |
| | Somatic Awareness | $7.63 \pm 2.54$ | $8.21 \pm 2.92$ |
| | Theory of Mind | $9.13 \pm 2.77$ | $10.04 \pm 1.95$ |
| | Validation | $14.24 \pm 2.53$ | $13.71 \pm 1.75$ |

*Note:* Values denote mean and standard deviation (SD). ASC = autism spectrum condition; CMP = comparison group.

**Table 4.**
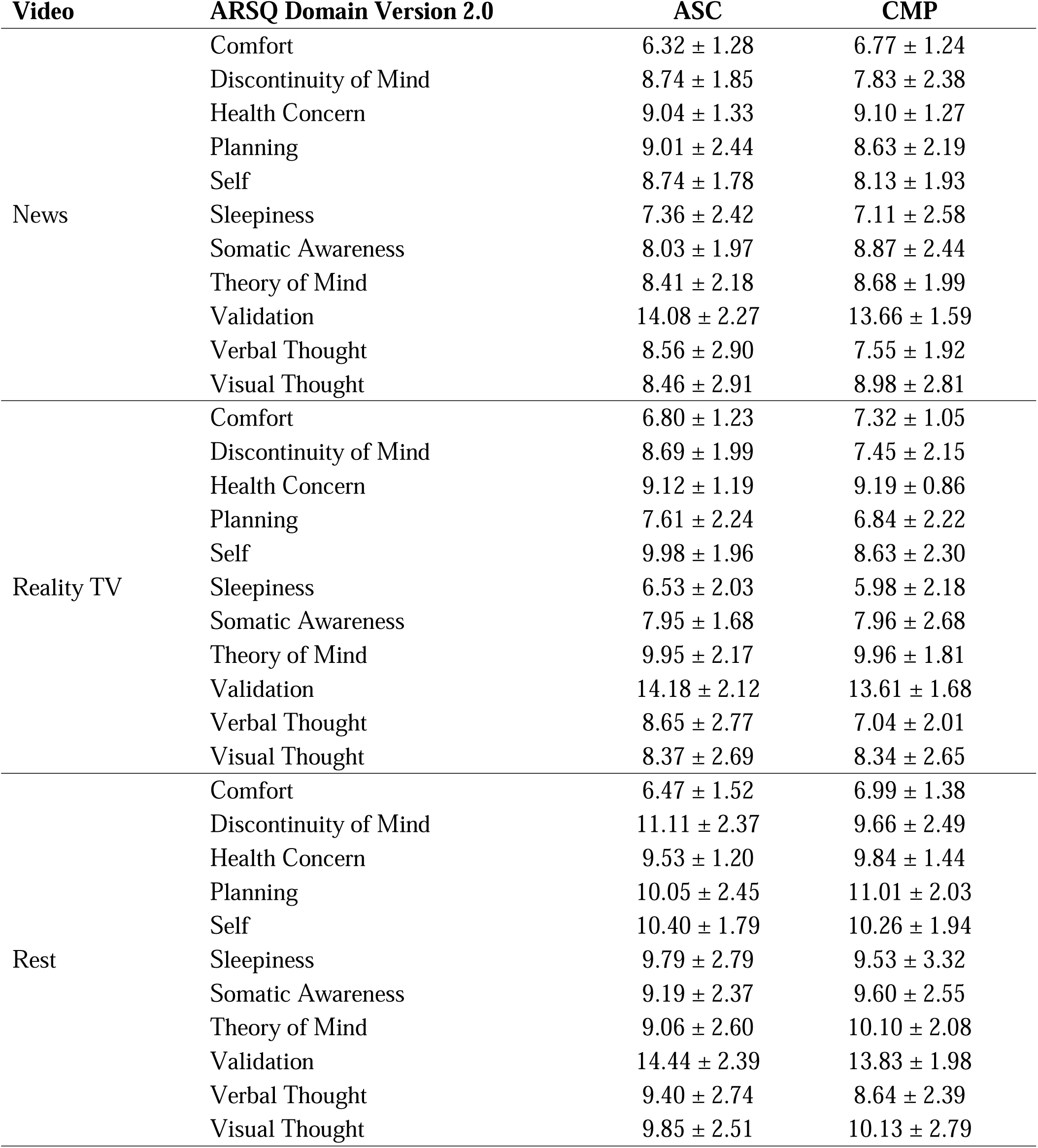
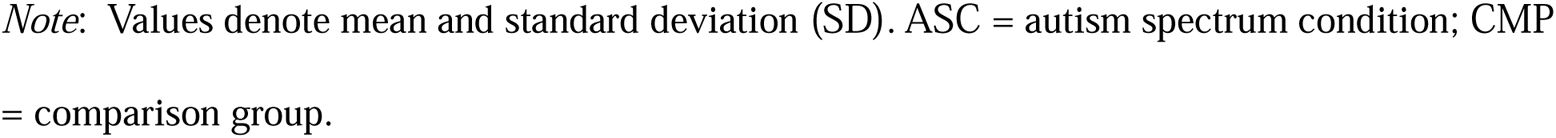
Descriptive statistics of Amsterdam Resting State Questionnaire (ARSQ) version 2.0.

| Video | ARSQ Domain Version 2.0 | ASC | CMP |
| --- | --- | --- | --- |
| News | Comfort | $6.32 \pm 1.28$ | $6.77 \pm 1.24$ |
| | Discontinuity of Mind | $8.74 \pm 1.85$ | $7.83 \pm 2.38$ |
| | Health Concern | $9.04 \pm 1.33$ | $9.10 \pm 1.27$ |
| | Planning | $9.01 \pm 2.44$ | $8.63 \pm 2.19$ |
| | Self | $8.74 \pm 1.78$ | $8.13 \pm 1.93$ |
| | Sleepiness | $7.36 \pm 2.42$ | $7.11 \pm 2.58$ |
| | Somatic Awareness | $8.03 \pm 1.97$ | $8.87 \pm 2.44$ |
| | Theory of Mind | $8.41 \pm 2.18$ | $8.68 \pm 1.99$ |
| | Validation | $14.08 \pm 2.27$ | $13.66 \pm 1.59$ |
| | Verbal Thought | $8.56 \pm 2.90$ | $7.55 \pm 1.92$ |
| | Visual Thought | $8.46 \pm 2.91$ | $8.98 \pm 2.81$ |
| Reality TV | Comfort | $6.80 \pm 1.23$ | $7.32 \pm 1.05$ |
| | Discontinuity of Mind | $8.69 \pm 1.99$ | $7.45 \pm 2.15$ |
| | Health Concern | $9.12 \pm 1.19$ | $9.19 \pm 0.86$ |
| | Planning | $7.61 \pm 2.24$ | $6.84 \pm 2.22$ |
| | Self | $9.98 \pm 1.96$ | $8.63 \pm 2.30$ |
| | Sleepiness | $6.53 \pm 2.03$ | $5.98 \pm 2.18$ |
| | Somatic Awareness | $7.95 \pm 1.68$ | $7.96 \pm 2.68$ |
| | Theory of Mind | $9.95 \pm 2.17$ | $9.96 \pm 1.81$ |
| | Validation | $14.18 \pm 2.12$ | $13.61 \pm 1.68$ |
| | Verbal Thought | $8.65 \pm 2.77$ | $7.04 \pm 2.01$ |
| | Visual Thought | $8.37 \pm 2.69$ | $8.34 \pm 2.65$ |
| Rest | Comfort | $6.47 \pm 1.52$ | $6.99 \pm 1.38$ |
| | Discontinuity of Mind | $11.11 \pm 2.37$ | $9.66 \pm 2.49$ |
| | Health Concern | $9.53 \pm 1.20$ | $9.84 \pm 1.44$ |
| | Planning | $10.05 \pm 2.45$ | $11.01 \pm 2.03$ |
| | Self | $10.40 \pm 1.79$ | $10.26 \pm 1.94$ |
| | Sleepiness | $9.79 \pm 2.79$ | $9.53 \pm 3.32$ |
| | Somatic Awareness | $9.19 \pm 2.37$ | $9.60 \pm 2.55$ |
| | Theory of Mind | $9.06 \pm 2.60$ | $10.10 \pm 2.08$ |
| | Validation | $14.44 \pm 2.39$ | $13.83 \pm 1.98$ |
| | Verbal Thought | $9.40 \pm 2.74$ | $8.64 \pm 2.39$ |
| | Visual Thought | $9.85 \pm 2.51$ | $10.13 \pm 2.79$ |

*Note:* Values denote mean and standard deviation (SD). ASC = autism spectrum condition; CMP = comparison group.

**Quality metrics for structural T1w & T2w scans.** The MRIQC pipeline v25.0.0 (Esteban et al., 2017) was used to extract image quality metrics of all T1-weighted (T1w) and T2-weighted (T2w) scans. When multiple T1w or T2w scans were available for the same participant, the highest quality scan was retained. To determine scan quality, four metrics were assessed: contrast-to-noise ratio (CNR), coefficient of joint variation (CJV), entropy-focused criterion (EFC) and ratio of median white-mater intensity to the 95th percentile of all signal intensities (WM2MAX). Because lower values of CJV and EFC indicate better quality, these metrics were inverted before summing the four measures to obtain a composite score; the scan with the highest composite score was retained for further analysis.

All metrics obtained for each participant and scan are reported in Table 8. Five metrics reflecting structural image quality are visualized in Figure 1. To determine whether quality metrics differed across participants with and without ASC, two-sample t-tests were performed on all quality metrics that showed variability across participants (Table 5). Overall, the ASC and comparison groups had largely comparable image quality for both T1w and T2w scans.

**Figure 1.**
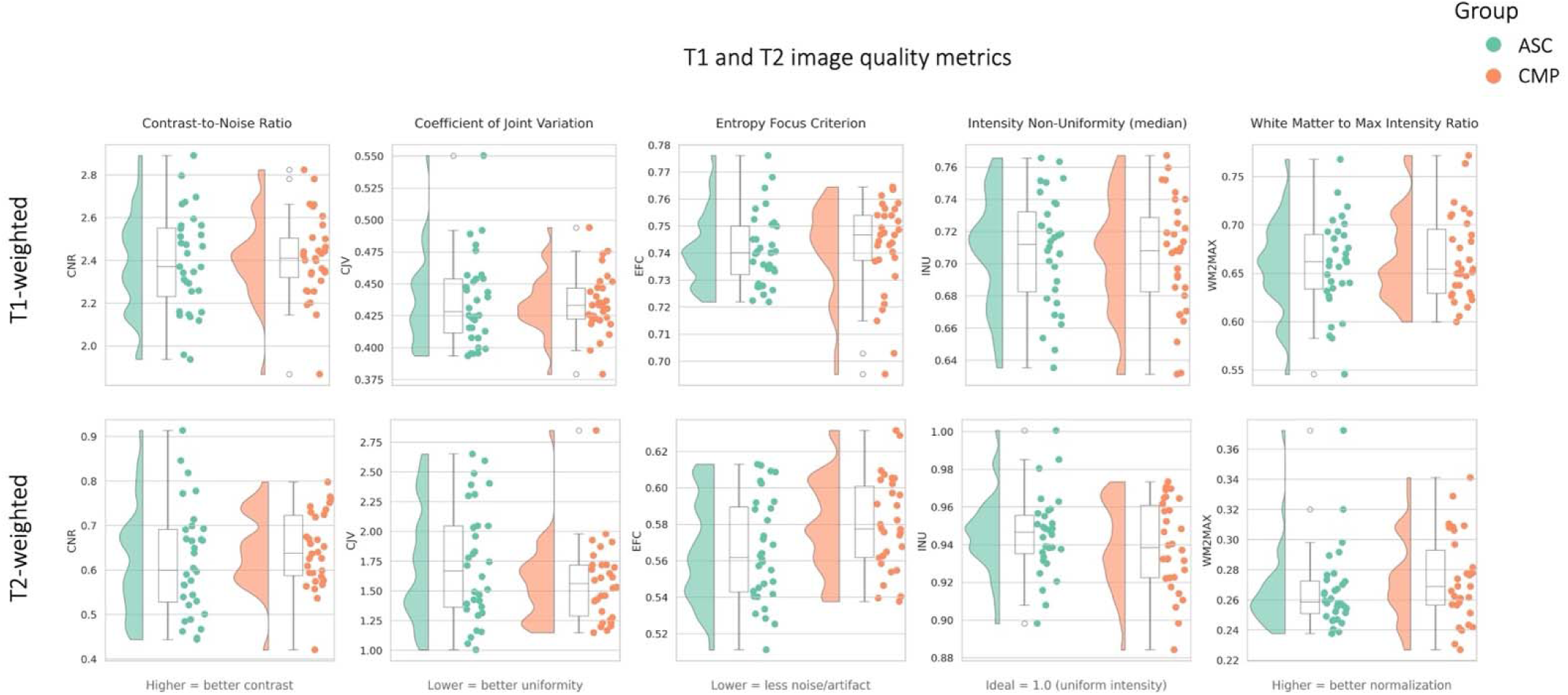
**Quality metrics related to T1-weighted and T2-weighted scans for autistic (ASC) and comparison (CMP) participants.** Contrast-to-Noise (CNR) ratio quantifies the contrast between grey and white matter; Coefficient of Joint Variation (CJV) reflects the variability between gray and white matter intensities and sensitivity to noise; Entropy Focus Criterion (EFC) indicates image sharpness and potential blurring or motion artefacts; median Intensity Non-Uniformity (INU) measures bias field inhomogeneity across the brain; white matter signal relative to maximum intensity (WM2MAX) reflects overall white matter signal relative to the image range.

For T1w images, participants with ASC showed higher spatial smoothness, as measured by full-width at half-maximum (FWHM) in the x and z directions and the average across all directions (all p<0.05). For T2w images, comparison participants had higher entropy-focused criterion (EFC) and a wider range of intensity non-uniformity (INU range) (all p<0.05). For all other metrics calculated by MRIQC there were no significant group differences.

Cortical surface reconstruction was performed using FreeSurfer v7.3.2 as part of fMRIPrep v25.2.3. Topological quality was assessed using the Euler characteristic, where values of 2 indicate topologically correct surfaces without defects. All 67 participants achieved Euler numbers of exactly 2.0 for both hemispheres. Grey-white matter contrast-to-noise ratio averaged 3.43 ± 0.09 (range: 2.97–3.62). CNR values were comparable between groups (ASC: 3.43 ± 0.08; CMP: 3.42 ± 0.11; t(65) = 0.58, p = 0.56).

**Quality metrics for functional scans.** The MRIQC pipeline was used to extract image quality metrics of all BOLD scans for each of the three runs per resting state condition. For each condition, quality metrics from all three runs were averaged within each diagnostic group. All metrics obtained for each participant and scan are reported in Table 9. Figure 2 displays four key metrics that reflect functional image quality. As was done for the structural data quality, two-sample t-tests were performed on all functional quality metrics that showed variability across participants to determine whether quality differed between groups (Table 6). Though groups differed across some quality metrics, ASC and comparison groups had comparable image quality for all fMRI conditions.

**Figure 2.**
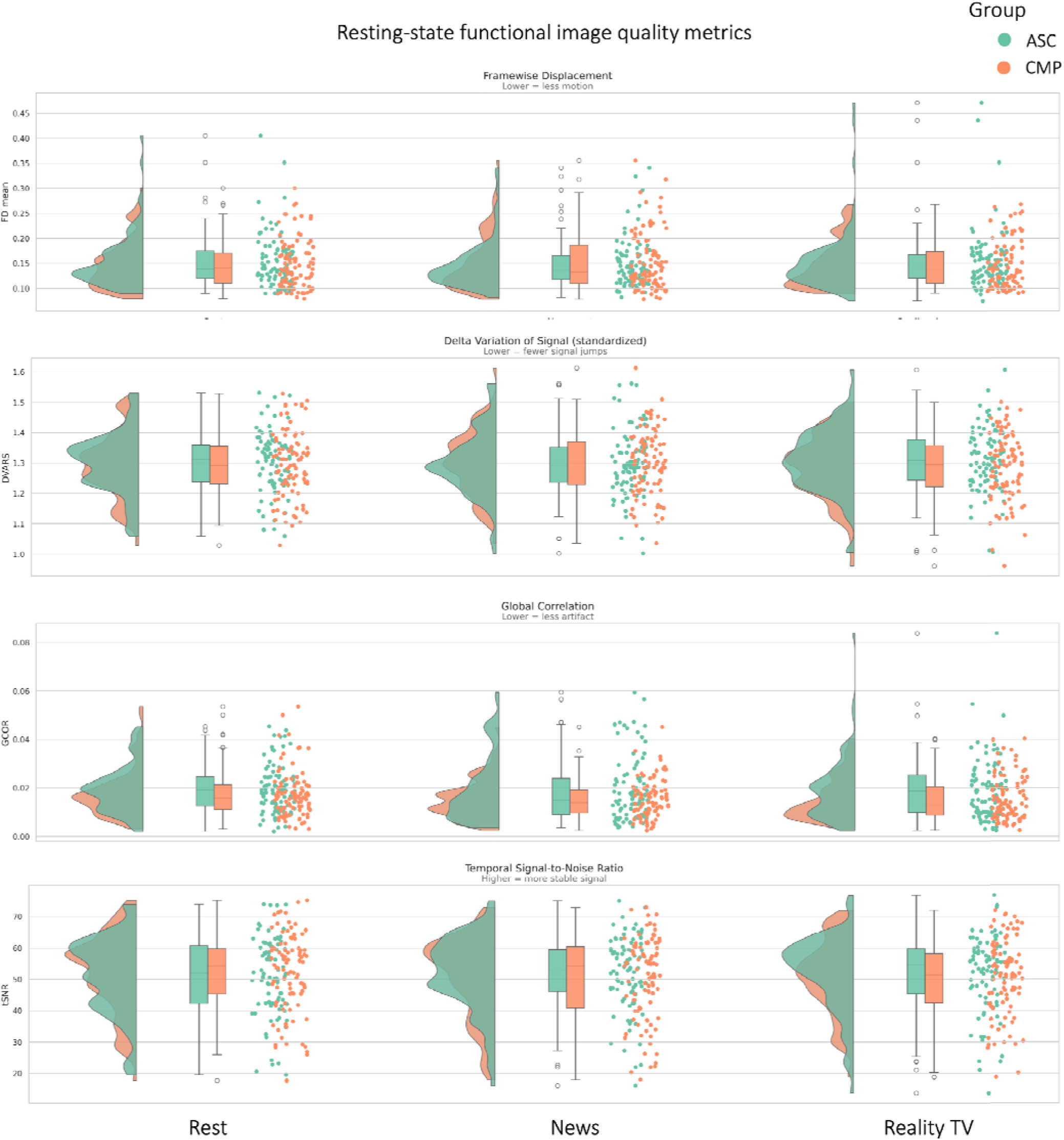
Quality metrics related to functional scans during three runs each of pure resting state, newscast viewing, and reality TV viewing for autistic (ASC) and comparison (CMP) participants. Framewise displacement (FD) quantifies the amount of participant head motion between consecutive volumes; standardized delta variation of signal (DVARS) quantifies the rate of change of the BOLD signal intensity between volumes, with high values indicating abrupt signal changes; global correlation (GCOR) measures the overall correlation across all brain voxels and is a reflection of global artefacts; temporal signal to noise ratio (tSNR) reflects the stability of the BOLD signal over time.

To further assess functional image quality, we calculated temporal signal-to-noise ratio (tSNR) maps by dividing the temporal mean by the temporal standard deviation at each voxel in the fMRIPrep-preprocessed data. Maps were calculated separately for each run and then averaged within participant before group averaging. Figure 3 shows group-averaged tSNR maps for both groups. As expected for gradient-echo echo-planar imaging (GE-EPI), tSNR values were highest in white matter (>100) due to minimal signal variability, whilst cortical grey matter showed tSNR values in the range of 50-100. Both groups demonstrated comparable spatial patterns of signal quality. There was signal dropout ventromedial prefrontal and anterior temporal regions, probably caused by susceptibility artefacts near air-tissue interfaces that affect gradient-echo sequences.

**Figure 3.**
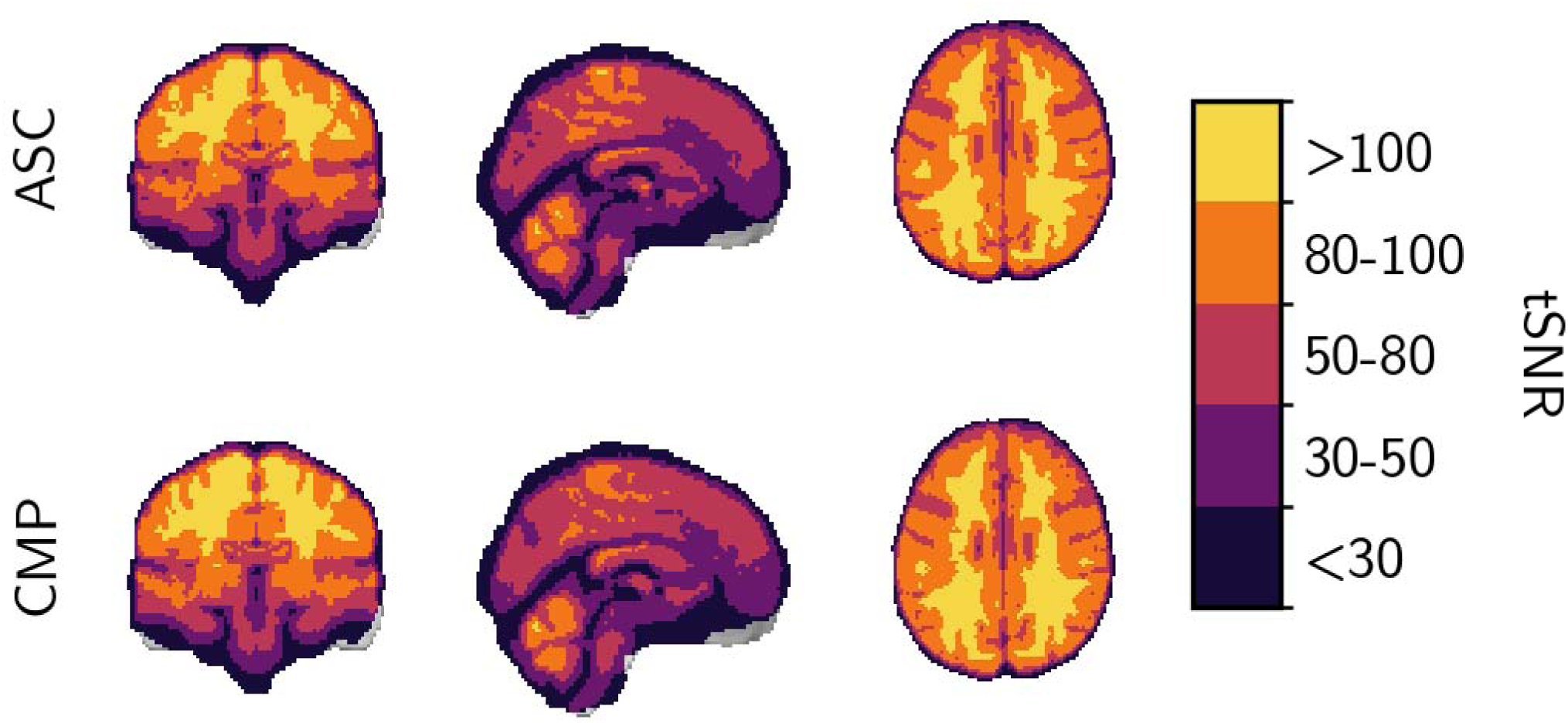
Group-averaged temporal signal-to-noise ratio maps for autistic (ASC) and comparison (CMP) groups. The tSNR maps indicate good functional data quality with tSNR >50 in cortical grey matter and characteristic signal dropout in ventromedial prefrontal and anterior temporal regions.

To validate data quality and participant engagement during the movie-watching conditions, we computed inter-subject correlation (ISC) maps using a leave-one-out approach, correlating each participant’s time series with the group average of all remaining participants. The data were parcellated using the Schaefer 400-parcel atlas. Strong ISC values in early sensory cortices (mean ISC in visual cortex = 0.410 ± 0.105, somatomotor cortex = 0.158 ± 0.091) confirm high signal quality and minimal noise artefacts. ISC values decreased systematically along the cortical hierarchy as expected, with more moderate correlations in association cortices (mean ISC = 0.215 ± 0.060). This pattern was broadly similar in the autism and comparison group (ASC: visual cortex mean ISC = 0.403 ± 0.101; association cortices mean ISC = 0.203 ± 0.060; CMP: visual cortex mean ISC = 0.416 ± 0.108, association cortices mean ISC = 0.226 ± 0.059), indicating equivalent data quality (Figure 4).

**Figure 4.**
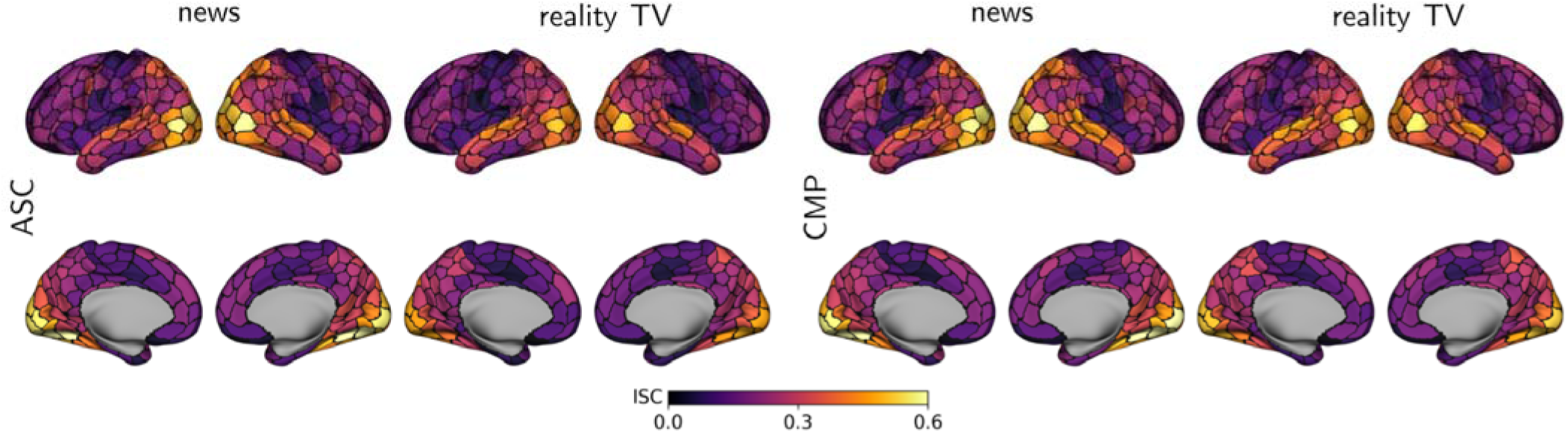
Group-average inter-subject correlation maps during naturalistic viewing. Inter-subject correlation (ISC) was computed using the leave-one-out approach. ISC maps were calculated separately for each clip and then averaged across the three clips within each stimulus category (news, reality TV). Maps are displayed separately for autistic (ASC) and comparison (CMP) participants. Both groups show highest ISC in early visual and auditory cortices, with additional engagement of parietal and anterior temporal association areas during both conditions. The colour scale indicates ISC values ranging from 0 (purple) to 0.6 (yellow).

Figure 5 displays average functional connectomes for the autism and comparison groups across the three fMRI conditions. Within each condition, the average connectivity patterns for the two groups are highly similar, indicating comparable data quality.

**Figure 5.**
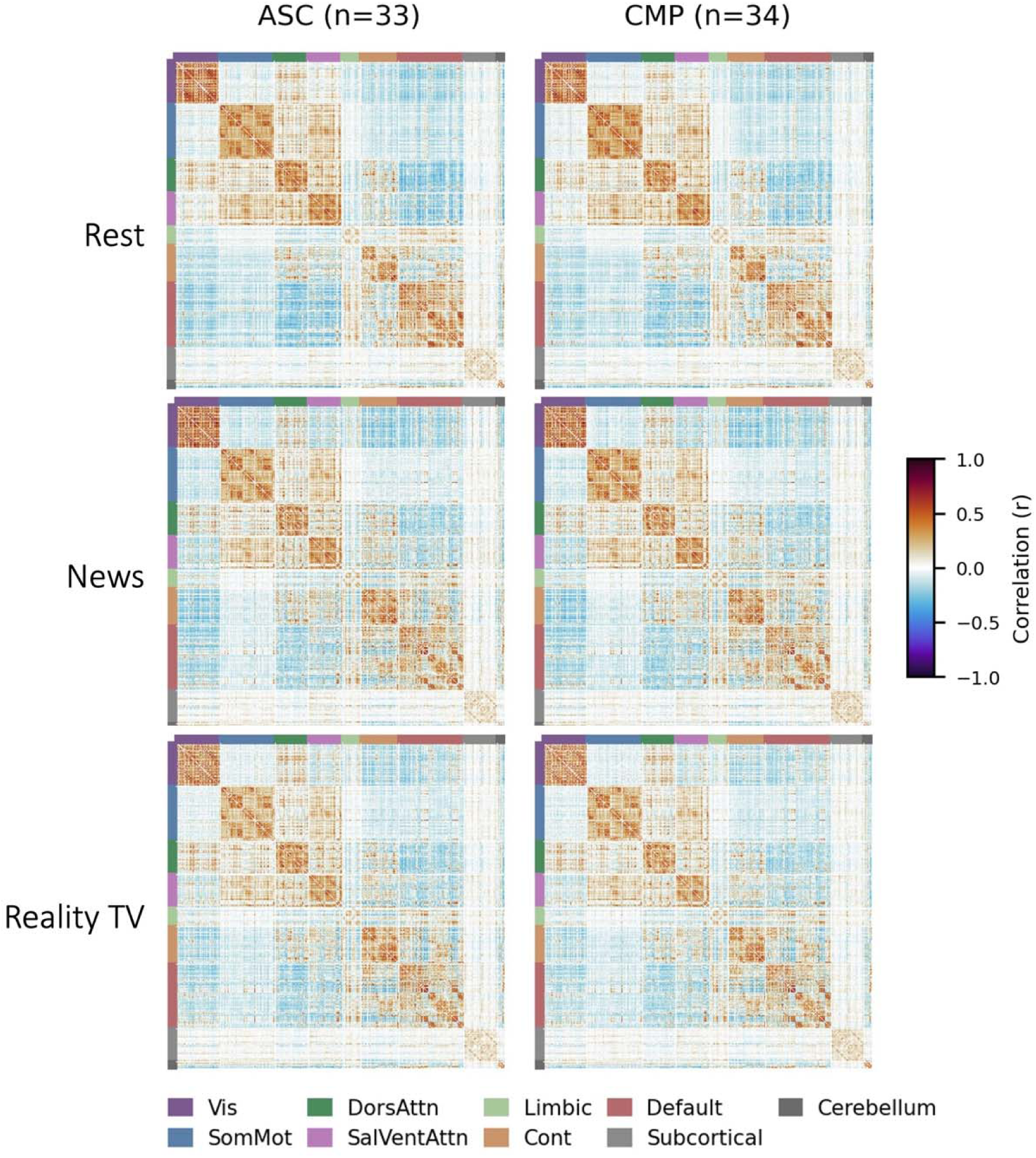
Group-averaged functional connectomes for autistic (ASC) and comparison (CMP) participants across three rsfMRI conditions. Group-level matrices were derived from relative-motion–corrected Pearson correlation matrices generated by XCP-D. Connectivity matrice reflect Pearson correlations between parcels from the Schaefer cortical atlas (parcellated at 400 parcels) supplemented with subcortical and cerebellar regions. Parcels are ordered by Yeo 7-network assignment, with white grid lines marking network boundaries. For each participant, functional connectivity matrices were computed separately for each run. The three runs belonging to the same condition (rest, news, or reality TV) were first averaged within subject, then averaged across participants within each diagnostic group. Both groups show highly similar connectivity patterns within condition, as well as strong within-network correlations, indicating comparable functional connectivity data quality across groups.

**Quality control for physiological data.** Physiological recordings (photoplethysmography for cardiac activity and respiratory belt for breathing) were collected simultaneously with fMRI acquisition. All 33 participants in the autism group and all 33 participants in the comparison group had physiological data available for at least one scanning session. Of 292 total fMRI sessions in the autism group, 274 (93.8%) had corresponding physiological recordings. In the comparison group, 282 of 298 (94.6%) fMRI sessions had physiological recordings. The missing recordings were due to technical issues or time constraints during specific scanning sessions.

Photoplethysmography (PPG) signal quality was assessed using the template-matching approach implemented in NeuroKit2 v0.2.12 (Makowski et al., 2021), which computes correlation coefficients between individual beat morphologies and an average template beat (Orphanidou et al., 2015). Recordings were classified as usable if >70% of samples achieved quality scores >0.5 and mean heart rate fell within the physiologically plausible range of 50-100 beats per minute.

Respiratory signal quality was assessed using both NeuroKit2’s automated quality metrics and manual inspection for extended flatline segments (>45 seconds), which indicate sensor disconnection. Recordings were deemed usable if they met the following criteria: (1) >70% of samples classified as good quality, (2) no extended flatline segments, (3) mean respiratory rate between 8-30 breaths per minute, and (4) minimum of 50 detected breaths per 10-minute recording. Brief periods of reduced or absent respiratory signal (10-30 seconds) were observed in some recordings and were retained as they likely reflect natural breath holds or very shallow breathing during focused attention, consistent with patterns reported in naturalistic viewing paradigms (Lynch et al., 2020). Only recordings with sustained technical failures (>45 seconds of continuous flatline or inability to detect respiratory cycles throughout the recording) were excluded.

Of the fMRI sessions with physiological data, quality control identified 267 of 274 (97.4%) cardiac recordings and all 274 (100%) respiratory recordings as usable in the autism group. In the comparison group, 270 of 282 (95.7%) cardiac recordings and 278 of 282 (98.6%) respiratory recordings met quality criteria. The small proportion of excluded recordings were due to sensor disconnection or technical failures during acquisition.

Mean heart rate was 71.5 ± 11.6 bpm in the autism group (n=267 recordings) and 69.5 ± 11.2 bpm in the comparison group (n=276 recordings). Mean respiratory rate was 17.4 ± 4.1 breaths/minute in the autism group (n=274 recordings) and 16.6 ± 3.7 breaths/minute in the comparison group (n=286 recordings). All values fall within expected physiological ranges for healthy young adults, confirming the biological validity of the recordings.

To assess the validity of physiological recordings, we examined the spatial distribution of variance explained by cardiac and respiratory regressors using a general linear model approach. Figure 6 displays group-averaged maps of absolute z-statistics for heart rate variability (HRV) and respiration volume per time (RVT) regressors across the autism (ASC) and comparison (CMP) groups. The spatial distribution of physiological effects followed anticipated patterns, with the strongest HRV effects in the brain stem and the strongest RVT effects in the white matter (Birn et al., 2006; Tong & Frederick, 2014). These results confirm that the acquired physiological recordings capture genuine cardiorespiratory fluctuations suitable for nuisance regression during functional connectivity analyses.

**Figure 6.**
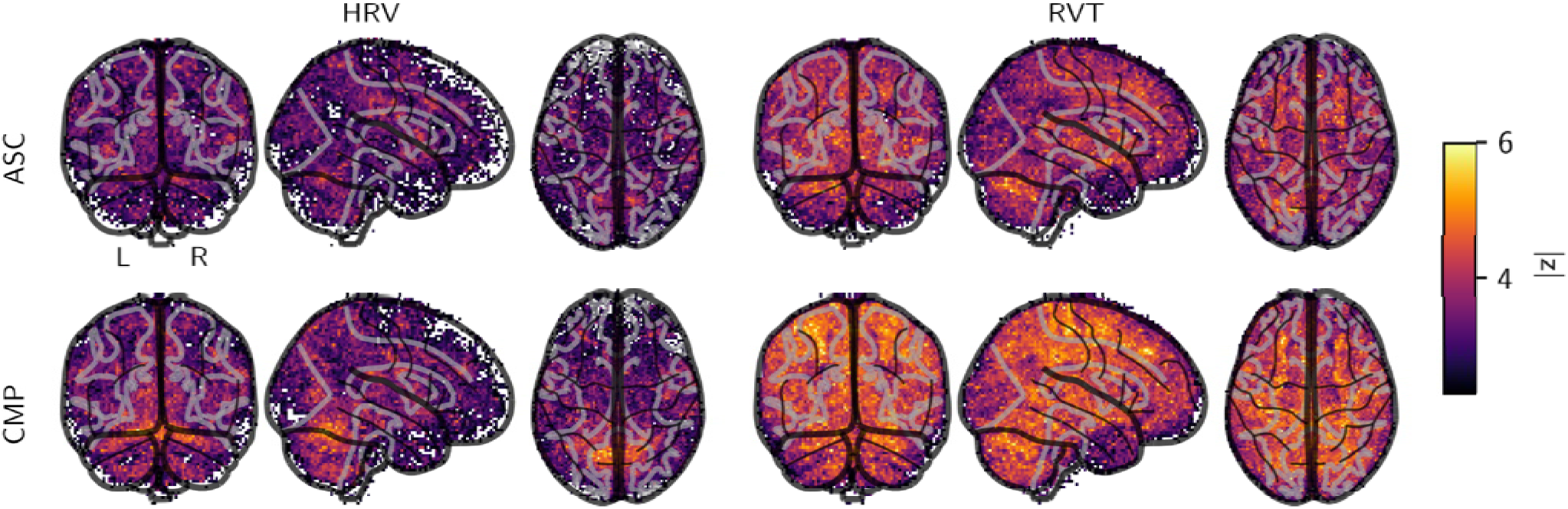
Spatial distribution of physiological noise effects for autistic (ASC) and comparison (CMP) participants. Group-level statistical maps showing absolute z-statistics for voxels where (left columns) heart rate variability (HRV) and (right columns) respiratory volume per time (RVT) regressors significantly explained BOLD signal variance. Top row: autism group (n=33); bottom row: comparison group (n=33). Maps thresholded at |z| > 2.3 (p < 0.01, uncorrected) and displayed on glass brain projections (coronal, sagittal, axial views). HRV effects were focal, primarily in deep grey matter, brainstem, and periventricular regions. RVT effects were widespread throughout cortical grey matter, with maximal effects in the ventricles.

**Quality metrics for diffusion-weighted scans.** We assessed diffusion MRI quality through visual inspection of group-averaged structural connectomes and fibre orientation maps. Probabilistic tractography was performed using QSIRecon v.1.1.1 with the mrtrix_singleshell_ss3t pipeline and anatomically constrained tractography. The group-averaged connectivity matrices demonstrate the expected hierarchical network organisation with stronger intra-network connectivity and symmetric homologous connections across hemispheres (Figure 7a). Direction-encoded colour maps reveal well-defined major white matter tracts including the corpus callosum, superior longitudinal fasciculus, and corticospinal tract (Figure 7b). Both groups show comparable structural connectivity patterns and fibre architecture, with no systematic artefacts or signal dropout. Quantitative quality metrics (Figure 8 and Table 7) confirmed that motion during acquisition was in the acceptable range (mean framewise displacement (FD [mean ± SD]): ASC (n=31): 0.4089 ± 0.0512 mm, range: 0.3246 - 0.4932 mm; CMP (n=33): 0.4054 ± 0.0420 mm, range: 0.3419 - 0.5060 mm).

**Figure 7.**
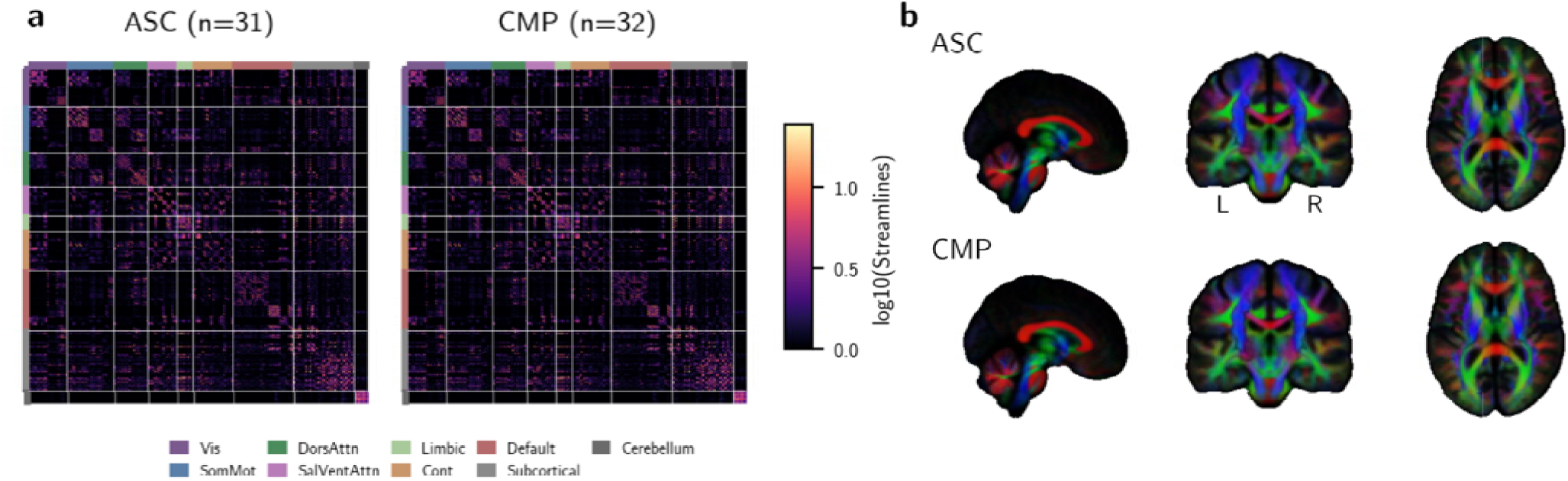
Diffusion MRI quality control for autistic (ASC) and comparison (CMP) groups. (a) Group-averaged structural connectomes derived from probabilistic tractography using *qsirecon*. Connectivity matrices show log -transformed streamline counts between regions of the Schaefer 200-parcel atlas, supplemented with subcortical and cerebellar parcellations. Parcels are ordered by Yeo 7-network assignment, with white grid lines delineating network boundaries. (b) Group-averaged direction-encoded color (DEC) maps showing principal fibre direction (red = left-right, green = anterior-posterior, blue = superior-inferior) modulated by apparent fibre density (AFD). Both groups show consistent connectivity patterns and expected white matter organisation, indicating comparable diffusion data quality.

**Figure 8.**
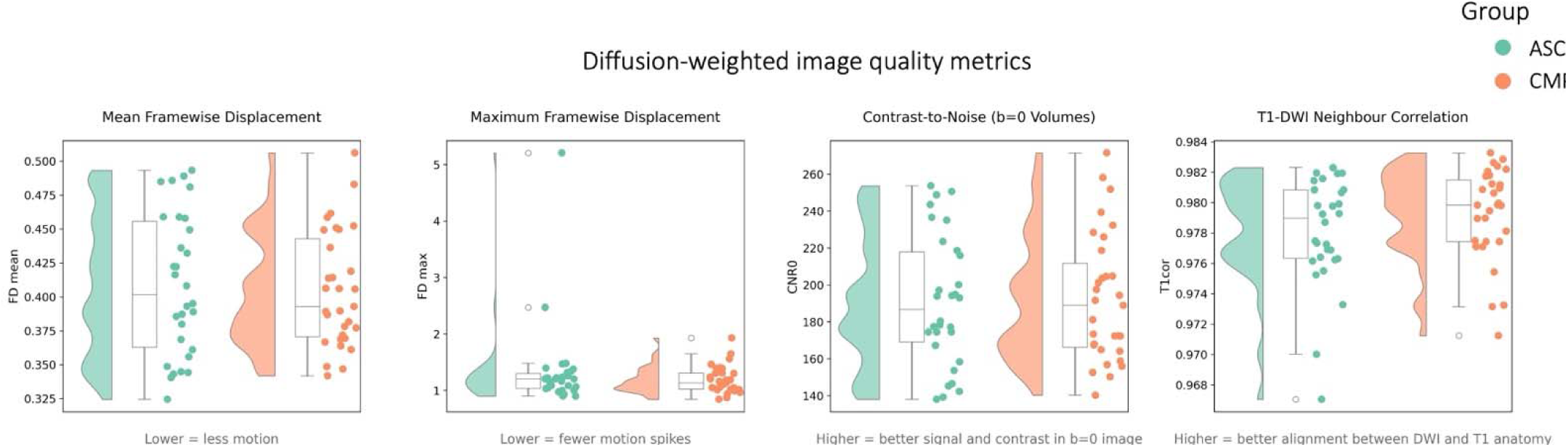
Quality metrics related to diffusion-weighted scans. Mean and maximum framewise displacement (FD) quantify the amount of participant head motion between consecutive volumes; Contrast-to-noise ratio in b=0 volumes (CNR0) reflects signal quality and image contrast; T1-DWI neighbor correlation (T1cor) measures alignment between the DWI scan and the participant’s T1-weighted anatomical scan.

## Usage Notes

The dataset and the stimulus materials are available at doi:10.18112/openneuro.ds007182.v1.0.0 and can be downloaded using the repository’s web interface or command-line tools. The complete raw dataset is approximately 133 GB; preprocessed derivatives add an additional 1.7 TB. Users may download specific modalities or individual participants as needed.

The dataset is provided in both raw and preprocessed formats. Raw data are suitable for users who wish to apply custom preprocessing pipelines or test novel methods. Preprocessed derivatives are provided for users prioritising analysis over preprocessing.

This dataset was designed for precision imaging analyses that prioritise within-individual characterisation over group averaging. With approximately 90 minutes of functional data per participant across three sessions, the dataset enables robust individual-level connectivity estimation, functional connectome fingerprinting, and person-specific parcellation. The three-session design supports test-retest reliability analyses, whilst the multiple viewing conditions (rest, news, reality TV) allow investigation of state-dependent network modulation within individuals. Users may also integrate this dataset with larger collections such as ABIDE or EU-AIMS LEAP through normative modelling or phenotypic harmonisation, as the dataset includes commonly used measures (AQ-60, WAIS-IV, SDQ).

Several important limitations should be noted. The single-shell diffusion acquisition (b=1000 s/mm², 128 directions) is suitable for DTI metrics and constrained spherical deconvolution-based tractography but does not support advanced multi-shell models such as NODDI or CHARMED. Gradient-echo echo-planar imaging exhibits characteristic susceptibility-induced signal dropout in ventromedial prefrontal and anterior temporal regions that cannot be corrected through preprocessing. Verbal ability assessment was changed after the first study phase from PPVT-NL to WAIS-IV Vocabulary due to ceiling effects. Therefore, assessments of verbal ability are not available for some participants, though Matrix Reasoning scores are available for all participants. Not all data are available for all sessions or imaging modalities; users should consult the data availability tables (Tables 10 and 11) and scans.tsv files to identify complete cases for specific analyses.

**Table 10.** Data availability of structural scans.

| **Structural scan** | **n** | **%** |
| --- | --- | --- |
| T1w | 66 | 100.0 |
| T2w | 63 | 95.5 |
| DWI | 61 | 92.4 |

**Table 11.** Data availability of functional scans, accompanying field maps, and physiological data from each condition per session.

|  | **Session 1** |  | **Session 2** |  | **Session 3** |  |
| --- | --- | --- | --- | --- | --- | --- |
| **Functional scan** | **n** | **%** | **n** | **%** | **n** | **%** |
| *Resting state* |  |  |  |  |  |  |
| BOLD image | 65 | 98.5 | 63 | 95.5 | 65 | 98.5 |
| Field map | 62 | 93.9 | 63 | 95.5 | 65 | 98.5 |
| Physiological | 61 | 92.4 | 59 | 89.4 | 60 | 90.9 |
| *Newsclip video* |  |  |  |  |  |  |
| BOLD image | 66 | 100.0 | 62 | 93.9 | 65 | 98.5 |
| Field map | 65 | 98.5 | 62 | 93.9 | 65 | 98.5 |
| Physiological | 62 | 93.9 | 58 | 87.9 | 63 | 95.5 |
| *Reality show video* |  |  |  |  |  |  |
| BOLD image | 66 | 100.0 | 64 | 97.0 | 65 | 98.5 |
| Field map | 65 | 98.5 | 64 | 97.0 | 65 | 98.5 |
| Physiological | 61 | 92.4 | 60 | 90.9 | 63 | 95.5 |

Users should cite this data descriptor when using the dataset and acknowledge the Dutch Research Council (NWO grant 406.XS.24.01.007) for funding the data collection. Questions about the dataset or analysis recommendations can be directed to.

## Code Availability

All code used to process and analyze this dataset can be retrieved from https://github.com/nadza-dz/PRISMA-Precision-Functional-Imaging-in-Autism.git.

## Author Contributions

ND: Data curation, Formal analysis, Methodology, Software, Validation, Visualization, Writing – original draft, Writing – review & editing

HG: Conceptualisation, Funding acquisition, Writing – review & editing HSS: Conceptualisation, Methodology, Validation, Writing – review & editing

JB: Conceptualisation, Data curation, Formal analysis, Methodology, Project administration, Software, Supervision, Validation, Visualization, Writing – original draft, Writing – review & editing

Author Note

## Acknowledgements

This publication is part of the project Precision Functional Imaging in Autism - A Comprehensive Study on Functional Brain Networks and State-Dependent Variability with file number 406.XS.24.01.007 which was partly financed by the Dutch Research Council (NWO).

We thank all participants for their time and contribution to this research. Further, we are grateful for the contributions of Fabiënne van Beek, Jette Benckhuisen, Juni Boumeester, Nando Braspenning, Noor Brink, Roos Heuschmidt, Helena Hoxha, Elisa Kanpaanpää, and Fleur van Pelt for their assistance with data collection.

---

Table 5 *Group comparisons of MRIQC-derived T1w and T2w image quality metrics between participants with and without autism spectrum disorder. (Available in supplementary excel document).*

Table 6 *Group comparisons of MRIQC-derived functional image quality metrics between participants with and without autism spectrum disorder. (Available in supplementary excel document).*

Table 7 *Group comparisons of QSIPrep-derived diffusion image quality metrics between participants with and without autism spectrum disorder. (Available in supplementary excel document).*

Table 8 *All image quality metrics related to T1-weighted and T2-weighted scans per participant derived from MRIQC. (Available in supplementary excel document).*

Table 9 *All image quality metrics related to functional scans per participant derived from MRIQC.* (Available in supplementary excel document).

